# Barcoding-inferred biodiversity of shallow-water Indo-Pacific demosponges

**DOI:** 10.1101/2024.12.29.630654

**Authors:** Dirk Erpenbeck, Adrian Galitz, Michael L. Berumen, Gabriele Büttner, Cécile Debitus, Moritz Dirnberger, Merrick Ekins, Kathryn Hall, Leonard Namuth, Sylvain Petek, Neda Rahnamae, Maggie M. Reddy, Daniela Rettenberger, Stefanie R. Ries, Simone Schätzle, Christine H.L. Schönberg, Edwin Setiawan, Joëlle van der Sprong, Olivier P. Thomas, Vani Tirumalasetty, Nicole J. de Voogd, Oliver Voigt, John N.A. Hooper, Gert Wörheide

## Abstract

**Aim:** The Indo-Pacific is the world’s largest marine biogeographic region. It is characterised by different degrees of connectivity among its subregions, and harbours the majority of demosponge species currently known to science. Comparisons between several regional sponge faunas have been undertaken in the past, mostly based on identifying the sponge species morphologically. The Sponge Barcoding Project, in tandem with other regional DNA taxonomy campaigns, provides one of the largest DNA-based taxonomic data collections from sponges of the Indo-Pacific. Here, we utilise the sponge barcoding data in the largest molecular biodiversity study of sponges to date, which reveals patterns of shallow-water demosponge faunal connectivity, endemism, and distribution in the Indo-Pacific with a level of resolution unavailable in prior morphology-based studies.

**Location:** Demosponge specimens in this study cover 13 marine provinces of the Indo-Pacific, from the Red Sea to South East Polynesia.

**Methods:** We classified demosponge barcodes using the ribosomal subunit (28S rDNA) of 1,910 sponge samples into molecular operational taxonomic units (MOTUs). MOTU composition of the 13 marine provinces was compared based on Jaccard and Sørenson dissimilarities, and other biodiversity indices.

**Results:** Our data corroborated high levels of endemism among demosponges. Faunal overlaps were revealed between the Red Sea and the Gulf, which displayed relatively small connectivity with other marine provinces of the Western Indian Ocean. In the Western Indian Ocean, we observed a strong faunistic boundary to the Central Indo-Pacific. The Polynesian sponge faunas were comparatively isolated marine provinces of the Central Indo-Pacific.

**Main conclusions:** Our data corroborate case studies on sponges that generally reject the presence of cosmopolitan or otherwise widespread sponge species, instead revealing high levels of regional endemism. This is consistent with similar observations and hypotheses in other marine invertebrates. Connectivity among Indo-Pacific marine provinces differs for demosponges in many aspects from that of other marine taxa, such as corals and fishes, probably due to their shorter pelagic larval phase.

## Introduction

The Indo-Pacific encompasses an expansive region of tropical and subtropical waters, extending from the East African coast, including the Red Sea in the west, to nearly 240 longitudes eastward beyond Easter Island. This vast area represents one of the most biodiverse marine regions globally (e.g., Stehli et al. 1967; Stehli and Wells 1971; Briggs 1987; Van Soest 1994). While the West Pacific is regarded as the marine region with the highest marine diversity worldwide, the Northwestern (Arabia) and Southwestern Indian Oceans constitute additional biodiversity hotspots, harbouring high proportions of endemic taxa (e.g., Hoareau et al. 2013; DiBattista et al. 2016a). Due to its extensive size, a comprehensive assessment of Indo-Pacific faunal zonations for understanding biogeography and dispersal patterns remains challenging.

The primary drivers of biogeographical differentiation are (1) the large distances of open ocean difficult to cross for species with limited dispersal capabilities, and (2) the oceanic currents determining the predominant directions of offspring distribution. Early studies of Ekman (e.g. 1953) and Briggs (e.g. 1974) defined marine biogeographic regions, realms, and provinces in the Indo-Pacific based on zoogeographic barriers and endemism, while the subdivision into marine ecoregions (Spalding et al. 2007) aimed to facilitate conservation efforts (Briggs and Bowen 2012). More taxon-based bioregionalisation models have been formulated for reef-fishes (Briggs and Bowen 2012; Kulbicki et al. 2013), corals (Keith et al. 2013; Veron et al. 2015), and other taxa. In this context, molecular studies in particular have highlighted faunal heterogeneity, frequently revealing that species previously thought to be widespread are, in fact, species complexes with allopatric “*mosaic*” distributions (e.g., Malay and Paulay 2010; DiBattista et al. 2012; Reid et al. 2013).

The Indo-Pacific is a biodiversity hotspot for Demospongiae, the largest sponge class that comprises 85% of all described sponge species (Briggs 1987; Hooper and Lévi 1994; Van Soest 1994). The assessment of overall sponge species richness is challenging (Hooper et al. 2013), and mostly restricted to smaller geographical studies with limited spatial or taxonomic range. While the Western Indian Ocean represents a significant “peak of species diversity” for several marine phyla (Obura 2016), it remains comparatively unexplored for sponges (Van Soest et al. 2012). Here, our current understanding of demosponge biodiversity is primarily based on species lists (e.g., Rützler 1972; Thomas 1979; Van Soest 1993, 1994), alongside few comparative studies (Barnes and Bell 2002).

Our knowledge of Western Indian Ocean sponges is less compared to other marine taxa, despite the increasing use of molecular markers (e.g. Ngwakum et al. 2021). Better assessments exist for adjacent regions, such as the Red Sea (see Wooster et al. 2019), the Gulf of Oman (Van Soest and Beglinger 2002, 2008), and are gaining momentum for the Eastern Arabian Sea (e.g., George et al. 2020), while the Persian Gulf sponge fauna remains underexplored (see Erpenbeck et al. 2020a). Comparatively more studies are available from the Malay Archipelago in terms of alpha and beta sponge diversity (e.g., Van Soest 1989, 1990; Bell et al. 2004, 2013; Cleary et al. 2005; Becking et al. 2006; de Voogd et al. 2006; de Voogd and Cleary 2008; Rovellini et al. 2019, and several others), although large parts of the Coral Triangle biodiversity hotspot (cf. Hoeksema 2007) are yet to be studied. Likewise, several thousand sponge species were reported for Australian waters, but a large proportion of them are new to science and awaiting description (e.g., Hooper and Lévi 1994; Hooper et al. 2002, 2013; Hooper and Ekins 2005; Fromont et al. 2016). Many other regions in the tropical Pacific were studied on a restricted spatial scale only (e.g. Lévi et al. 1998; Kelly et al. 2003, 2016; Hall et al. 2013; Freeman and Easson 2016; Núñez Pons et al. 2017), lack comprehensive data (Van Soest et al. 2012), or were only recently the subject for broader scale biogeographic analyses (Galitz et al. 2023, 2024). Similarly, most of these studies lack data on cryptobiotic sponges, which represent a considerable portion of the taxonomic diversity on coral reefs (Richter et al. 2001; Timmers et al. 2020).

Our limited understanding of sponge diversity stands in sharp contrast to their immense ecological importance. Especially on coral reefs, sponges take on key functions in several facets: they protect exposed areas of coral skeletons from boring organisms through overgrowth, stabilise coral fragments in disturbed reefs and thus facilitate reef consolidation, and contribute significantly to reef bioerosion through the activity of excavating sponges (Bell 2008; Wulff 2016). Of particular importance for the oligotrophic coral reef habitats, is the role of sponges in the conversion of compounds from dissolved to particulate, or *vice versa* (Hammond and Wilkinson 1985; Corredor et al. 1988; de Goeij et al. 2013). Knowledge of sponge biodiversity and biogeography is indispensable for the recognition of faunal changes, the application of conservation measures, or even the implementation of a “blue economy” (Kelsh et al. 2011; Glover et al. 2018).

Unfortunately, reliable and comparable assessment of sponge species richness has proven difficult, particularly when using classical (*i*.*e*., morphology-based) identification techniques (Hooper and Lévi 1994; Van Soest et al. 2012; Hooper et al. 2013). Shortcomings are due to the paucity, homoplasy, and plasticity of characters, and challenges of taxonomy and phylogenetic reconstruction (e.g., Fromont and Bergquist 1990; Erpenbeck et al. 2006; Cárdenas et al. 2011). Likewise, the lack of an objective species concept for sponges hampers the classification into meaningful units for faunal comparisons (see e.g. Setiawan et al. 2016 for a summary). In the last decades, DNA sequence-based methods provided biodiversity assessment tools requiring comparatively little expertise in local faunas, while offering greater objectivity in species identification. The Sponge Barcoding Project was established as a centralised platform to gather molecular taxonomic data on sponge species worldwide, in order to facilitate unambiguous identification methods (Wörheide and Erpenbeck 2007). Here, we utilise sponge barcoding data to investigate patterns of sponge faunal connectivity across the Indo-Pacific. We aim to gain insight into endemism and species distribution of sponge faunas of marine provinces from the Red Sea to eastern Polynesia. Additionally, this largest molecular biodiversity analysis, to date, seeks to identify dispersal boundaries for sponges in the Indo-Pacific in comparison to other marine taxa.

## Material and Methods

### Origin of data

The basis of the current analysis is shallow water (mostly SCUBA collected) material from various biodiversity collections stretching from the far Western (Red Sea, Mayotte) to the Eastern (Polynesia) Indo-Pacific. Barcoding data were generated in the course of the Sponge Barcoding Project (Wörheide and Erpenbeck 2007) or during independent, ongoing, barcoding studies. All sequences are retrievable from the European Nucleotide Archive (see Appendix S1), metadata of specimens sequenced in the course of the Sponge Barcoding Project are available in the Sponge Barcoding Database (www.spongebarcoding.org).

All specimens were allocated to their respective marine provinces (MP) following Spalding *et al*. (2007 i.e., “distinct biotas that have at least some cohesion over evolutionary time frames”). These MPs were used as a geographic baseline for biodiversity comparison. Data from the Red Sea (MP: *Red Sea and Gulf of Aden*) originated from biodiversity surveys along the Saudi Arabian coastline conducted 2012-2014 (published in Erpenbeck et al. 2016). Persian Gulf samples (MP: *Somali/Arabian*) were collected during several campaigns in Iran using SCUBA diving from 2015 to 2017 (Erpenbeck et al. 2020a). Maldives (MP: *Central Indian Ocean Island*s) samples were obtained during fieldwork in 2017 and 2022 in Magoodhoo (Fafuu Atoll). Mayotte (MP: *Western Indian Ocean*) samples were collected during the ANR-Netbiome project in May 2013) and retrieved from the collections of Naturalis Biodiversity Center (Leiden, the Netherlands). Eastern Indo-Pacific samples originated from French Polynesia (MP: *South East Polynesia*) and Wallis Island (MP: *Central Polynesia*), and were recently published by Galitz *et al*. (2023, 2024). Samples from MPs *Northeast Australian Shelf*, *East Central Australian Shelf*, *Tropical Southwestern Pacific*, and most samples from MP *Sahul Shelf* constituted collection material from the Queensland Museum, Brisbane, and the Australian Institute of Marine Science, and were added to the Sponge Barcoding Project (Erpenbeck et al. 2012, 2020b; Vargas et al. 2012), as was comparative material from MP *Southeast Australian Shelf*. Sequence data from Papua New Guinea (MP *Eastern Coral Triangle*) were provided for this project from yet unpublished biodiversity surveys. Material for MP *Sunda Shelf* was assembled from the collections of the Naturalis Biodiversity Center and the Bavarian State Collection for Paleontology and Geology Munich (SNSB-BSPG).

### Sequence generation and alignment

Analyses were based on the C-region of the nuclear large ribosomal subunit (28S), a ∼450 bp standard marker for sponge DNA barcoding and biodiversity assessments (Voigt and Wörheide 2016). DNA was extracted following the plate extraction method developed for sponge barcoding (Vargas et al. 2012), by spin columns or using CTAB (Sambrook and Russell 2001). Subsequently, 28S-C region fragments were amplified, purified, and both strands sequenced following previously published protocols (Erpenbeck et al. 2016). All assemblies were manually checked using CodonCodeAligner v3.7 (www.codondode.com). Positions where potential Intergenomic Polymorphisms (IGPs) were detected as characteristic double peaks in forward and reverse sequences were coded with the respective IUPAC ambiguity codes, which resulted in their exclusion during the subsequent MOTU clustering (see below).

### Molecular OTU (MOTU) delineation

Sequences were divided with respect to their respective marine provinces *sensu* Spalding *et al*. (2007) using QGIS 3.10. (QGIS Development QGIS Development Team 2019, http://www.qgis.osgeo.org). Sequences longer than 340 bp were aligned with ClustalW (Thompson et al. 1994) as incorporated in the msa package for R (Bodenhofer et al. 2015), which facilitates an unambiguous alignment of sequences with low genetic distances in a data set consisting of highly variable sequences from a wide phylogenetic range. In this approach, the performance of ClustalW was insensitive to differences in sequence length and presence of ambiguity codes. DECIPHER 2.0 (Wright 2016) was used to cluster sequences into the respective MOTUs using the unweighted pair group method with arithmetic mean (UPGMA) algorithm (following Cowman et al. 2017; Hadiyanto et al. 2021), which has been shown to have the best performance in hierarchical cluster analyses (Kreft and Jetz 2010).

We chose a stringent MOTU cutoff of 0.3%, which was found to consider the genetic differences detected in several case studies between selected sympatric shallow water demosponge species (e.g., Erpenbeck et al. 2017, 2020b; Ekins et al. 2023), and was subsequently used in smaller-spatial biodiversity studies (Galitz et al. 2023, 2024). As this stringency demands high fidelity of the sequencing, no external sequences (*e*.*g*., NCBI Genbank sequences published by other authors) have been used, as their correct basecalling could not be verified by us. As an additional quality test, we repeated the biodiversity analyses, restricting MOTUs consisting of a minimum of two sequences only (*i*.*e*., excluding singletons) to rule out bias from undetected sequencing or basecalling errors.

### Biodiversity analyses

We compared the different MPs in terms of their beta diversity, and the measurement of species composition difference between species assemblages. Rarefaction analyses on species richness and the sampling completeness of the respective marine provinces were conducted with iNEXT (Chao et al. 2016). Biodiversity analyses were performed with the ‘picante’ (Kembel et al. 2010) and ‘vegan’ (Oksanen et al. 2013) packages for R (see Appendix S2 for results). As sampling has been predominantly qualitative, *i*.*e*. without measurement of abundance data, the only visible differences between sites are in species identities (Barwell et al. 2015).

From the large number of beta-biodiversity indices for absence-presence data (see Koleff et al. 2003), we used the Jaccard index as it is comparatively invulnerable to errors of taxonomy and enumeration, and has relatively low error rates for geographic undersampling (Jaccard 1901). In parallel, the Sørensen index was calculated (Sørensen 1948); the Sørensen index is closely related to the Jaccard index, but gives double the weight to the shared species, and therefore reduces the errors caused by false negatives (Pos et al. 2014; Schroeder and Jenkins 2018; Hadiyanto et al. 2021). The significance of the beta diversity indices was subsequently verified with PERMANOVA (permutational multivariate analysis of variance) using a custom variation of the *adonis* function of ‘vegan’ for pairwise comparison of regions, with 999 permutations and adjustments for false discovery rate (FDR) taken into account (Anderson 2001, 2017). Visualisation of MOTU overlap was conducted using UpSetR (Conway et al. 2017). The cryptobenthic fauna of coral reefs, however, has a different taxonomic composition, and has been disregarded in this analysis for this reason (Timmers et al. 2020; Vicente et al. 2021). Likewise, other factors potentially affecting the taxonomic composition, such as temporal variability in sponges could not be considered, but it is not anticipated that this will bias the overall taxonomic pattern (see e.g., Beepat et al. 2020).

## Results

A total of 1910 sequences of 28S data from 12 marine provinces was analysed (see Table 1). The 28S barcoding region is of variable length and had an average of 386 basepairs (bp) in our data set (median 388 bp, maximum 481 bp; short fragments are due to restrictive clipping in the case of ambiguous basecalling at the 5’ or 3’ terminus). After restriction to a sequence length > 340 bp, the data set comprised 1,823 sequences (average 402 bp, median 399 bp). Sequence numbers per region varied between 306 (MPs *Red Sea and Gulf of Aden*) and 57 (*Tropical Southwestern Pacific*). These sequences fell into 701 MOTUs with the maximum again from MP *Red Sea and Gulf of Aden* (126), and a minimum from MP *Somali / Arabian* (31). A total of 414 MOTUs (59.1%) were singletons, consisting of one specimen only in the entire collection. Taxonomically, the 701 MOTUs are dominated by 169 Dictyoceratida (24.1%) and 144 Haplosclerida (20.5 %); all other orders were less abundant.

**Table 1:** Overview of regional sequence yield and MOTU numbers in the current analysis. * overall number of different MOTUs among all marine provinces in this analysis

| Marine Province | Seqs. total | Seqs. included | MOTUs | MOTUs (excl. singlet.) | endemic MOTUs | endemic MOTUs (excl. singlet.) |
| --- | --- | --- | --- | --- | --- | --- |
| ***Central Polynesia*** | 302 | 297 | 105 | 57 | 75.2% | 54.4% |
| ***Central Indian Ocean Islands*** | 280 | 267 | 71 | 44 | 76.1% | 64.4% |
| ***East Central Australian Shelf*** | 79 | 73 | 45 | 27 | 53.3% | 22.2% |
| ***Eastern Coral Triangle*** | 84 | 78 | 50 | 24 | 70% | 37.5% |
| ***Somali/Arabian*** | 73 | 73 | 31 | 18 | 67.7% | 44.4% |
| ***Northeast Australian Shelf*** | 207 | 185 | 99 | 56 | 59.6% | 28.6% |
| ***Red Sea and Gulf of Aden*** | 303 | 299 | 126 | 52 | 84.1% | 61.5% |
| ***Sahul Shelf*** | 166 | 154 | 97 | 39 | 77.3% | 43.6% |
| ***Southeast Polynesia*** | 133 | 130 | 89 | 38 | 78.7% | 50.0% |
| ***Sunda Shelf*** | 130 | 126 | 34 | 24 | 67.6% | 54.2% |
| ***Tropical Southwestern Pacific*** | 57 | 47 | 39 | 20 | 56.4% | 15.0% |
| ***Western Indian Ocean*** | 96 | 94 | 51 | 24 | 72.5% | 41.7% |
| ***Total*** | **1910** | **1823** | **701\*** | **287\*** | **Ø 70,43%** | **Ø 43,12%** |

Comparison between MOTU distribution indicated a high percentage of MOTUs (86.3%) restricted to a single marine province (= endemic MOTUs, see Table 1). Levels of endemism ranged from 53.3% (MP *East Central Australian Shelf*) up to 84.1% (MP *Red Sea and Gulf of Aden*), with an average of 70.43% per MP. Quantification of the shared MOTUs of every MP can be deduced from Figure 1.

**Figure 1:**
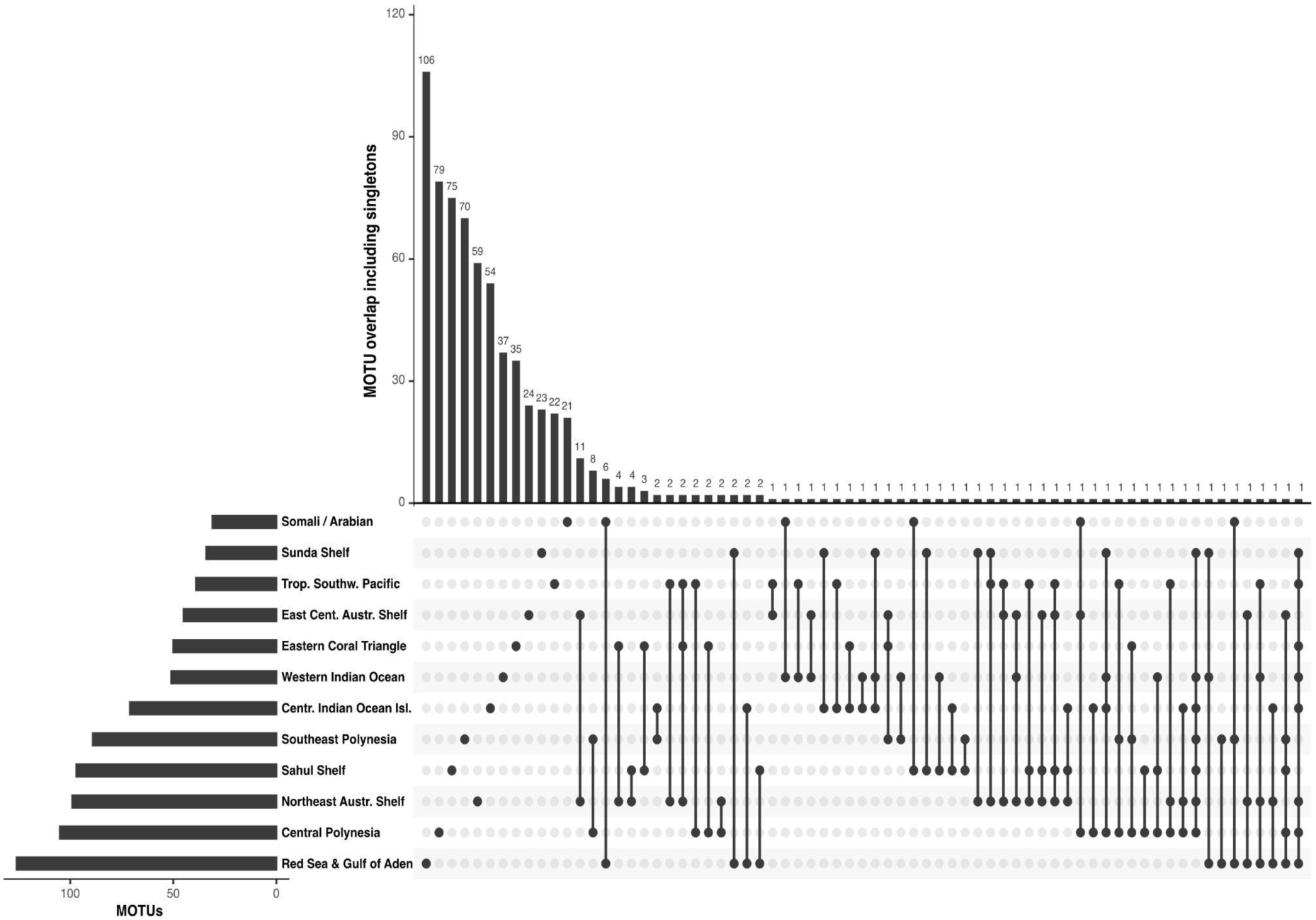
MOTU count including singletons per marine province and overlap between provinces. Histogram on the left: Number of MOTUs per MP. To its right: Black dots without connecting lines represent the number of unique MOTUs in a MP, while dots connected by lines indicate the number of MOTUs present in two or more MPs. Counts of MOTUs for each scenario depicted with histogram on the top.

Jaccard dissimilarity indices between the marine provinces recover MP pairs without MOTU overlap (=dissimilarity of 100) and those with MOTU overlaps to different degrees (Figure 2). Jaccard dissimilarities were also plotted on a map of the Indo-Pacific (Figure 3). We recovered several distinct MP clusters, three of them with Jackknife support ≥ 60: i) *Central Polynesia* with *South East Polynesia*; ii) *Central Indian Ocean Islands* with *Western Indian Ocean* and *Sunda Shelf*; iii) *Somali/Arabian* with *Red Sea and Gulf of Aden*; iv) *East Central Australian Shelf* with *Northeast Australian Shelf* and (to a lesser extend) *Tropical Southwestern Pacific*. Analyses excluding singletons, or based on Sørensen indices (including and excluding singletons), revealed congruent results that differed only in the clustering of *Sahul Shelf* or the *Eastern Coral Triangle* with provinces of the latter clade (Appendix S3, Figs S3.1-S3.3).

**Figure 2:**
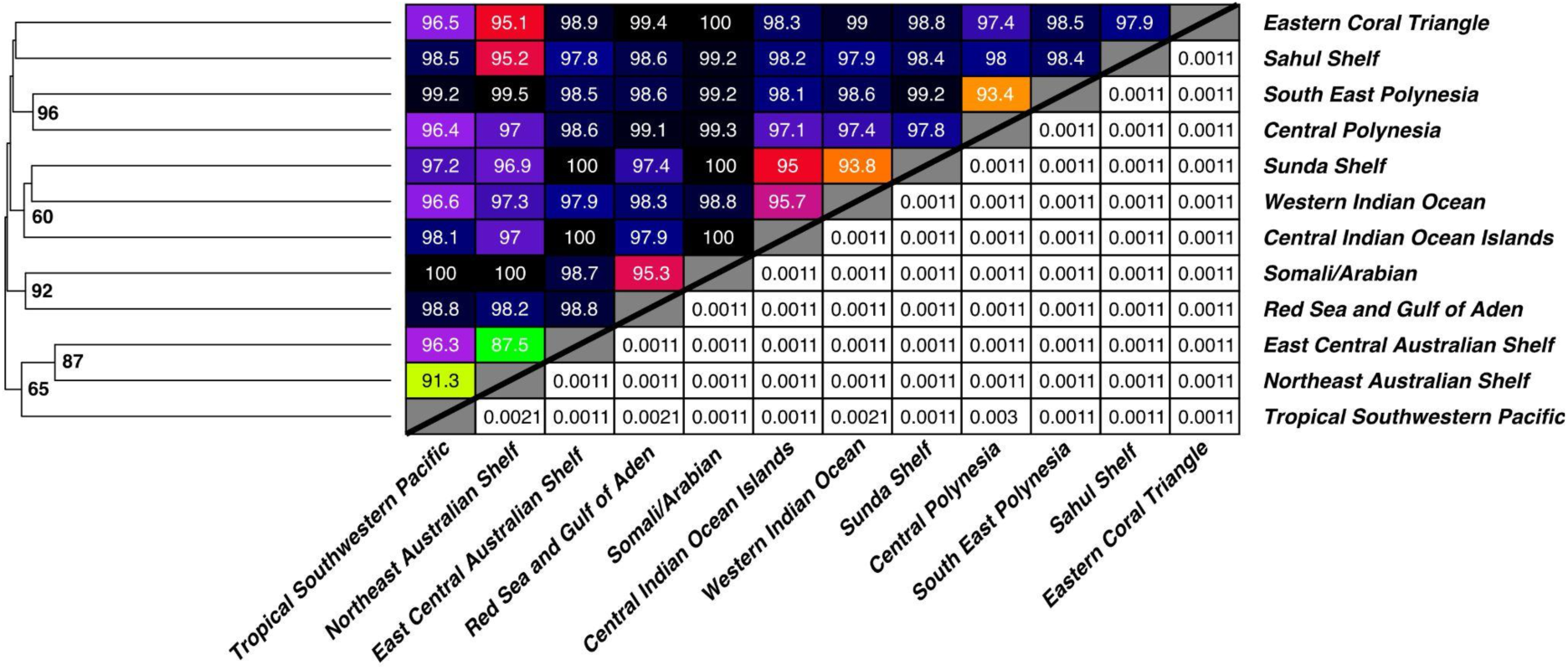
Diagram top left: 28S Jaccard dissimilarities of the studied marine provinces. Cell colour corresponds to the legend depicted in Figure 3 plus black in case of complete dissimilarity (100). Diagram bottom right: False discovery rate (FDR) adjusted p values of respective 28S Jaccard indices generated through pairwise PERMANOVA. The dendrogram on the left depicts faunal similarities between two given regions based on their respective Jaccard indices. Numbers on nodes are for Jackknife support (1000 replicates).

**Figure 3:**
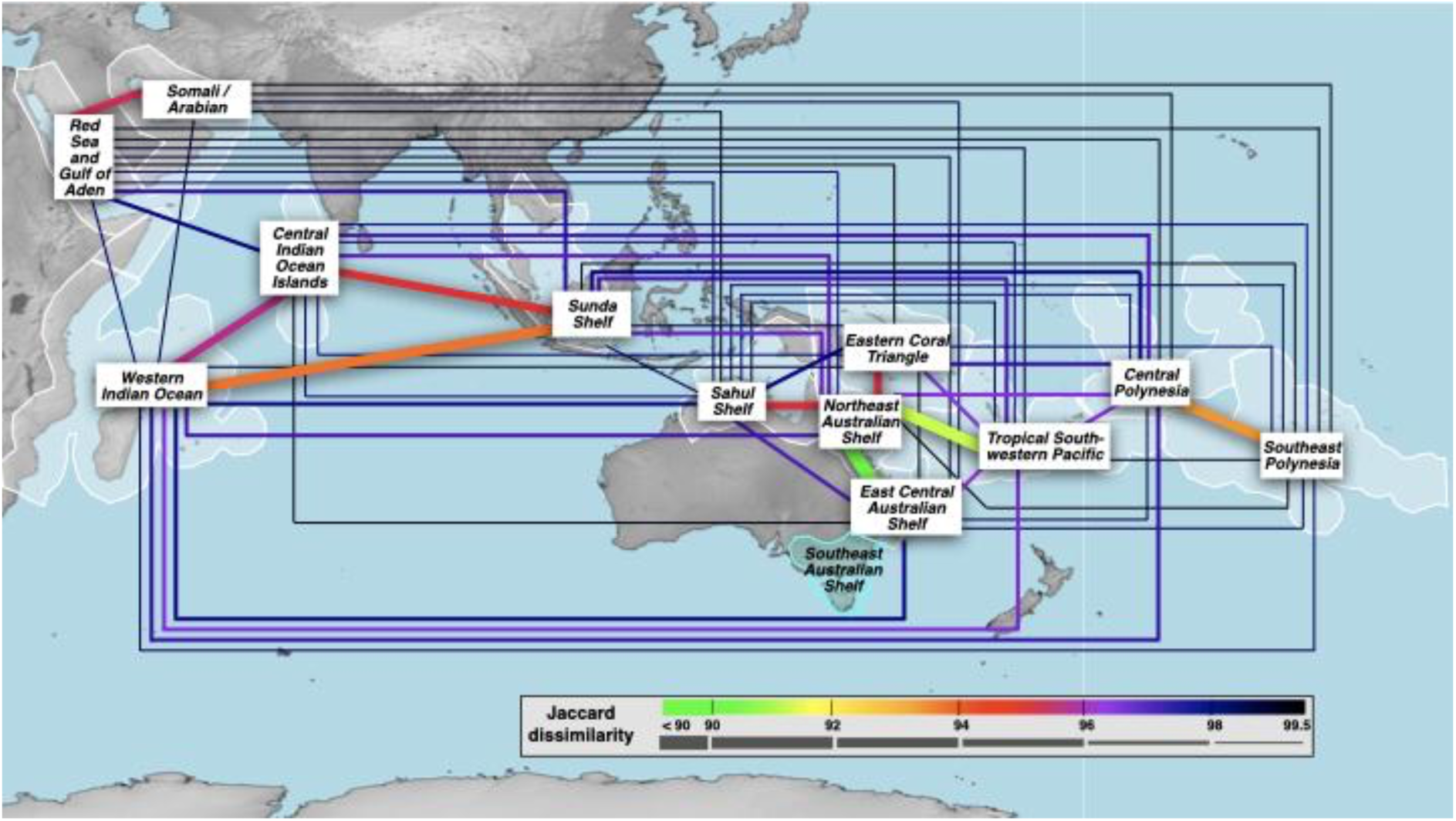
Jaccard dissimilarities between the samples of the twelve Indo-Pacific marine provinces, including singletons. Tropical and subtropical marine province areas and boundaries are shaded in white, temperate in light blue. Colour and thickness of the connecting lines between tropical and subtropical marine provinces indicate their dissimilarity levels. Colours are similar to Figure 1. Jaccard dissimilarities of 100 (i.e. no shared MOTUs) are not indicated by a line. Jaccard dissimilarities between 99.5 and 99.9 were not present. Map generated with SimpleMappr (Shorthouse 2010).

## Discussion

### High demosponge endemism due to limited dispersal potential

Our study, the largest molecular analysis on shallow water demosponge biodiversity to date, shows a high percentage of non-overlapping MOTUs between the marine provinces, indicating a high degree of endemism throughout the Indo-Pacific region. This pattern parallels earlier molecular studies that repeatedly rejected the frequent occurrence of widespread sponge species (e.g., Wörheide et al. 2002; Fromont et al. 2006; Pöppe et al. 2010; Reveillaud et al. 2010; Xavier et al. 2010). Our results are therefore consistent with earlier hypotheses, which projected a generally high rate of endemism among other marine invertebrates (e.g., Palumbi et al. 1997; Klautau et al. 1999; Solé-Cava and Boury-Esnault 1999; Barnes and Bell 2002; Bierne et al. 2003). In this respect, several genetic studies on sponges and other marine organisms have found genetic structure at large spatial scales, suggesting that long-distance panmixia and dispersal are rare (e.g., Lazoski et al. 2001; Whalan et al. 2008; Benestan et al. 2021), with only few demosponge species currently being discussed as widespread (e.g., Carballo et al. 2013), or even globally-distributed (e.g., Turner 2020). In congruence with this, our study identified only few MOTUs occurring in the western *and* eastern extensions of the analysed Indo-Pacific MPs. The most frequently occurring MOTUs are MOTU 214 (in 8 of the 12 MPs), and MOTU 79 (in 7 MPs, see Fig.1 and Appendix S1). Both MOTUs have previously been identified as *Stylissa carteri* (Dendy 1889) and “*Hyrtios* cf. *erectus*”, with the former including specimens frequently misidentified as *S. massa* (Carter 1887), and the latter being a cryptic, but widespread sister lineage of *H. erectus* Keller (1889) (Erpenbeck et al. 2017). Our data underline that, in contrast to most other MOTUs, both taxa are highly common in the Indo-Pacific (see de Voogd and Cleary 2008).

Crandall *et al*. (2019) argued that physical or environmental barriers play a limited role in the wide, spatial genetic differentiation of the Indo-Pacific. Therefore, dispersal capabilities for larvae constitute a significant factor for the biodiversity patterns of demosponges (Maldonado 2006). In many shallow-water species the larvae is a planktonic lecithotrophic parenchymella with a short life-span of a few days only, during which time a suitable substrate for the sessile adult stage must be occupied (unlike the hoplitomella of some deeper water tetractinellids that can survive several weeks, Maldonado 2006). Here, the absence of suitable stepping stones in the vast oceanic distances reduce distribution success for sponges, resulting in pronounced range fragmentation (see Mora et al. 2012). Other demosponge lineages possess benthic, rather than planktonic larvae, which crawl on the substrate for several days and consequently possess even more limited dispersal capacities (Maldonado 2006). Passive means of transport were conclusively reported in only a few instances, *e*.*g*., by ballast water (Dunstan and Bax 2008), hull fouling (Carlton and Fowler 2018), or attachment to debris (Elvin et al. 2018; Al-Khayat et al. 2021). Propagules, clonal fragments, or enduring bodies (e.g., Leong and Pawlik 2010) have been described from only a few marine species, and should contribute little to demosponge dispersal processes in the Indo-Pacific.

### Demosponge connectivity in the Northwestern Indian Ocean

We found MOTUs overlapping between the Red Sea (MP *Red Sea and Gulf of Aden*) and the Persian Gulf (MP S*omali/Arabian*). Both water bodies constitute northern marginal seas of the Indian Ocean, and have shared physical similarities. Both are semi-enclosed seas in arid regions with scarce freshwater influx and high evaporation (Ludt et al. 2017), potentially favouring settlement and growth of similar demosponge species, as is found for reef-building corals (Obura 2012; Berumen et al. 2019). However, both seas have only a narrow and particularly shallow connection to the Arabian Sea, hindering the exchange of species already adapted to moderate depths (Ludt et al. 2017; see also Wang et al. 2019). Likewise, cold upwellings in the northern Arabian Sea form a barrier for temperature-sensitive species, and prevent a connection of coral reef stepping-stones to the other marine provinces (e.g. Klausewitz 1989; Kemp 1998; Siddall et al. 2003; Bailey 2010; DiBattista et al. 2016b), resulting in the high degree of demosponge endemism as observed previously (Erpenbeck et al. 2020a) and in the present study.

### Low connectivity between Northwestern and Western Indian Ocean

The demosponge distributions, as recovered from our molecular analyses, indicated a distinct pattern of considerably few MOTU overlaps between the northwestern Indian Ocean MPs (*Somali/Arabian* and *Red Sea and Gulf of Aden*) and the MPs of the remaining Indian Ocean. The MPs *Central Indian Ocean Islands*, *Western Indian Ocean*, and *Sunda Shelf* formed a triangle with a higher internal MOTU overlap isolated from the northwestern Indian Ocean MPs. Our findings matched earlier faunistic analyses on sponges of the Indian Ocean that favoured East - West connectivity of sponges rather than North - South (e.g., Indonesia to East Africa, see Van Soest 1994; Van Soest et al. 2012; Erpenbeck et al. 2017). This boundary divides the tectonically inactive western Indian Ocean and the active northern Indian Ocean seas. The cold upwelling currents in the Gulf of Aden and the Somali coast prevent the formation of shallow-water coral reefs as stepping-stones for sponges along the coast, and there is limited other habitat suitable for larval settlement alongside the Somali coast (Kemp 1998; DiBattista et al. 2013). The sponge larvae would have to cover vast distances in the open ocean to reach other MPs (e.g., the distance between Socotra near the Gulf of Aden to the Maldives, southwest of India is ∼2,000 km). In most cases these distances exceed the survival capabilities of the sponge larvae. For corals, which frequently possess a considerably longer pelagic larval duration than sponges (Nozawa and Okubo 2011; Miller et al. 2020), the northern Indian Ocean Gyre and the reversing Somali Current are important dispersal vectors between the northwestern and the remainder of the Indian Ocean (Obura 2012). These current systems appear less significant for the distribution of sponges.

We observed MOTU overlaps between the MPs *Central Indian Ocean Islands*, *Western Indian Ocean*, and *Sunda Shelf*, indicating a degree of connectivity between far western and eastern Indian Ocean MPs. Exchange is facilitated by the stepping stones of the Chagos Archipelago (called “Chagos Stricture” for corals, see Veron 1995) in combination with the South Equatorial Current and the Equatorial Countercurrent, rather than north-southernly currents. We did not find a division between Western Indo-Pacific and Central Indo-Pacific faunas, contrary to Spalding *et al*. (2007, based on multiple taxa and biophysical conditions), Kulbicki *et al*. (2013, reef fish), Briggs & Bowen (2012, 2013, based mostly on reef fish), or a similar boundary suggested for corals (Obura 2012, 2016; Veron et al. 2015). In multispecies analyses, no boundary in that region has been found to be significantly different from random distributions either (Crandall et al. 2019). Instead, more demosponge MOTUs of MP *Sunda Shelf* are shared with the *Western Indo-Pacific* realm than with other MPs of its *Central Indo-Pacific* realm. In this respect, demosponge MOTU distribution follows the Indo-Pacific Barrier along the western edge of the Sunda Shelf, which strengthened during low sea-level stands at the Last Glacial Maximum of the Pleistocene (Benzie 1999; Ludt and Rocha 2015; Crandall et al. 2019).

Several coral reef organisms of the MP *Sunda Shelf* have also been found to conform to the genetically differentiated “Western Province’’ of the Indo-Malay Archipelago (to the west of Makassar Strait), which has more resemblance to Indian Ocean faunas than to those of the Central Indo-Malay Archipelago (see Ducret et al. 2022 and references within). These findings are consistent with our observation that MP *Sunda Shelf* demosponges resemble (Western) Indian Ocean faunas more closely than any of the geographically closer MPs further east examined in this study.

### A marine “Wallace’s Line” for demosponges

Our data suggest a deep split of demosponge MOTU overlaps between MP *Sunda Shelf* and *Sahul Shelf* (including all other MPs of the *Central Indo-Pacific* and *Eastern Indo-Pacific* realms), despite their spatial proximity. This separation is congruent with cluster analyses of species lists by Van Soest *et al*. (2012). The Makassar Strait is a remnant of the biogeographic separation of Indian and Pacific Ocean marine populations during the Pleistocene emergence of the Sunda and Sahul shelves (occasionally termed “marine Wallace’s Line”, Barber et al. 2000; Reid et al. 2013), and marks a boundary. The current genetic distinctiveness of marine taxa still reflects the historical separation of Indian and Pacific Ocean basins, which was physically far greater during the glacial maxima (Barber et al. 2000). Several currents of the Indonesian Throughflow (ITF) contribute to the present separation. They pass through the Makassar and the Lifamatola Passage, eventually forming the South Equatorial or Leeuwin Current further west, therefore hampering larval and propagule exchange between habitats on both sides of the ITF (e.g., Kochzius and Nuryanto 2008; Reid et al. 2013).

### High connectivity within the Central Indo-Pacific Realm …and beyond

Our data found a comparatively high overlap of MOTUs between MPs *East Central Australian Shelf*, *Northeast Australian Shelf*, *Sahul Shelf*, *Tropical Southwestern Pacific*, and to a lesser extent, the *Eastern Coral Triangle*. Despite being a biodiversity hotspot, the *Eastern Coral Triangle* was included here in a comparative sponge biodiversity analysis for the first time (see Van Soest et al. 2012). The observed MOTU overlaps were a result of both spatial proximity and the presence of numerous stepping stones in this area that are connected by the East Australian Current.

Sponge species richness and taxonomic distributions of Australia have been the subject of several studies (e.g., Hooper 1994; Hooper et al. 2002; Hooper and Ekins 2005). These identified a major faunal transition zone at Cape York, which separates the MP *Sahul Shelf* at the Gulf of Carpentaria from the adjacent eastern MPs *Northeast Australian Shelf* (from Torres Strait to the Great Barrier Reef). Hooper and Ekins (2005) discussed this boundary as ecologically (habitat) related rather than biogeographic. Our present analysis corroborated this boundary, showing a higher Jaccard dissimilarity of MP *Northeast Australian Shelf* to MP *Sahul Shelf* than to MP *Tropical Southwestern Pacific*, which comprises the Coral Sea habitats on the Queensland Plateau and western Pacific Islands adjacent to the Great Barrier Reef.

Our data also observed a high MOTU overlap between MP *Northeast Australian Shelf* and MP *East Central Australian Shelf* to its south, displaying the lowest Jaccard dissimilarity between MPs in our analysis. Both MPs, however, are classified as different marine realms (realm *Central Indo Pacific* and realm *Temperate Australasia* respectively; (Spalding et al. 2007). We therefore expect a comparatively high Jaccard dissimilarity between these MPs, at least higher than between MPs of the same realm. However, Hooper and Ekins (2005) previously noted that a “well-recognised biogeographic transition zone” for sponges exists, along with a subsequent hard faunistic boundary to the temperate regions, but located further south of the current *Central Indo Pacific* realm limits. For this reason, we included the available barcoding data of the more southerly MP *Southeast Australian Shelf*, which is classified in realm *Temperate Australasia*, similarly to MP *East Central Australian Shelf* (Appendix S4, Fig. S4.1). We recovered a comparatively strong Jaccard dissimilarity of >0.98 between MP *East Central Australian Shelf* to MP *Southeast Australian Shelf*, while the dissimilarity to MP *Northeastern Australian Shelf* in the north was considerably lower (0.88) and among the lowest dissimilarities observed in this study (Appendix S4, Fig. S4.2). The results of our molecular genetic analyses therefore corroborated the findings of Hooper and Ekins (2005), indicating a strong faunistic boundary separating tropical from temperate faunas, located further south than suggested in the realm definitions of Spalding *et al*. (2007).

### Eastern Indo-Pacific in “splendid isolation”?

In the Eastern Pacific coral reefs, the abundance of macroscopic epibenthic sponges has been discussed as gradually decreasing in eastward direction due to different primary productivity requirements in the oligotrophic waters (Wilkinson 1987; Van Soest et al. 2012; Vicente et al. 2021). Sponge molecular biogeographic connectivity in that region has recently been analysed by Galitz *et al*. (2023), whose data are incorporated here to compare the *Eastern Indo-Pacific* realm MPs (MPs *Central Polynesia* and *Southeast Polynesia*) with the other Indo-Pacific MPs in the study. Our data revealed that sponge faunas of both MPs of the *Eastern Central Pacific* Realm are more similar to each other than to any MP of the *Central Indo Pacific*, or any other realm. This aligns with the hierarchical classification of Spalding *et al*. (2007), and corroborates the findings of Galitz *et al*. (2023) with a larger data set. Nevertheless, MP *Central Polynesia* still displayed a considerable degree of overlap with adjacent MPs of the neighbouring *Central Indo-Pacific* realm, which characterises MP *Central Polynesia* as a “melting pot of biodiversity” Galitz *et al*. (2023).

Crandall *et al*. (2019) identified spatial distance as the predominant factor reducing genetic exchange in the South Pacific, and only to a lesser extent the geologic history (continental and uplifted islands of Central Indo-Pacific realm and volcanic islands of the Eastern Indo-Pacific realm respectively, see Vermeij 1987). Here, the numerous small gyres of the South Pacific Current facilitate connectivity between demosponge faunas of Fiji and Tonga (MP *Tropical South Pacific*) with Wallis, Futuna, and Samoa (MP *Central Polynesia*) across the realm boundaries (see Galitz *et al*. 2023 for further details). Drivers of genetic exchange within the Eastern Indo-Pacific realm MPs are the South Pacific Current and South Equatorial Current. However, the vast distances (> 3,000 km) in combination with the short pelagic larval duration of most shallow-water demosponges hampers a higher MOTU overlap between its MP *Central Polynesia* and MP *Southeast Polynesia*.

## Acknowledgements

Parts of the subsampling was facilitated by the Marine Barcode of Life initiative (MarBol), funded by the Alfred P. Sloan Foundation. The scientific research cooperation between King Abdulaziz University (KAU), Faculty of Marine Sciences (FMS), Jeddah, Saudi Arabia, and the Senckenberg Research Institute (SRI), Frankfurt, Germany, in the framework of the Red Sea Biodiversity Project, during which the present material was collected, was funded by KAU GRANT NO. “I/1/432-DSR”. The authors acknowledge, with thanks, KAU and SRI for technical and financial support. Additional Red Sea fieldwork was supported by the King Abdullah University of Science and Technology (Award No. CRG-1-2012-BER-002 and baseline research funds to M.L.B) with special thanks to members of the Reef Ecology Lab for field assistance. Samples from Maldives were received with help and support from staff and scientists of MaRHE Center of University of Milano-Bicocca, Magoodhoo, and local collaborators (Agreement (AGR)438-ENV/PRIV/2024/83 incl. enclosed permits). DE is grateful for financial support from Lehre@LMU, ES for “WCP like 2023” funds of ITS Global Engagement Surabaya to DE. The Mayotte fieldwork was financed through the ANR-Netbiome under grant No ANR-11-EBIM-0006. Research permits were issued via Terres Australes en Antartiques françaises (TAAF). We thank Anne Gauvin-Bialecki, Bruno Fichou, Stephan Aubert, Philippe Prost, and Jean-Pierre Bellanger for their support.

## Notes

### Competing Interest Statement

The authors have declared no competing interest.

