## Appendix S2 for "Barcoding-inferred biodiversity of shallow-water Indo-Pacific demosponges"

**Figure S2:** Left panels: Rarefaction and extrapolation of biodiversity for each marine province analysed in the present MS based on sample-size; Right panels: Sample completeness for each marine province analysed in the present MS.

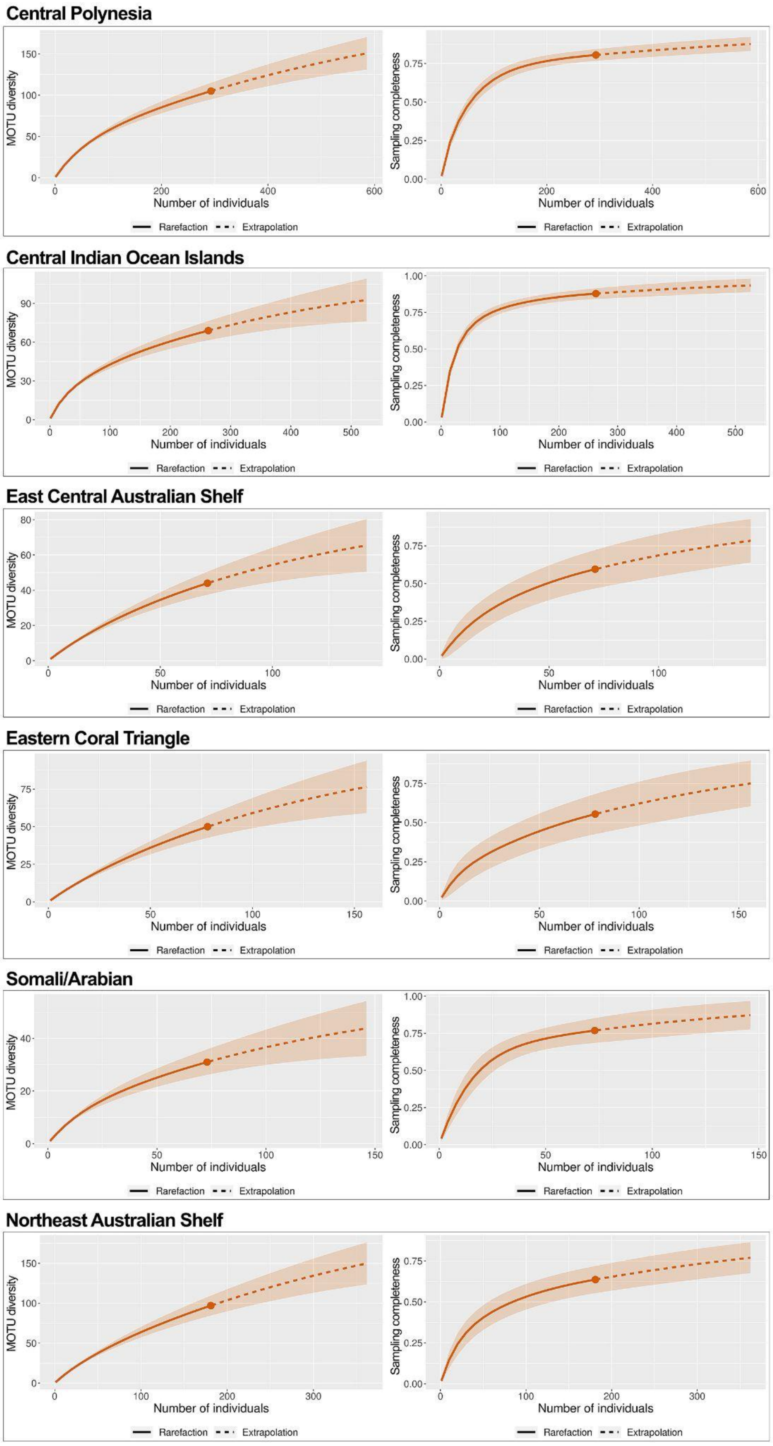

### Appendix S2 (continued)

#### Red Sea and Gulf of Aden

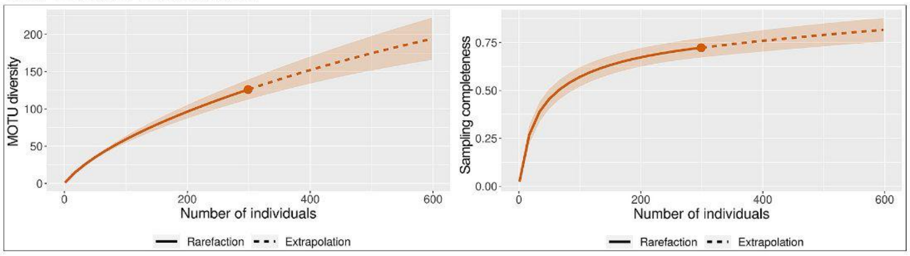


#### Sahul Shelf

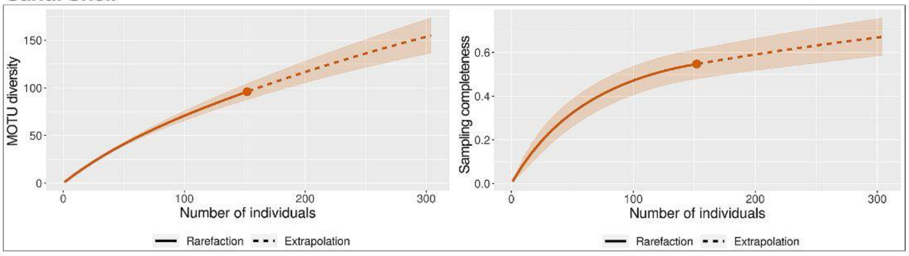


#### Southeast Polyensia

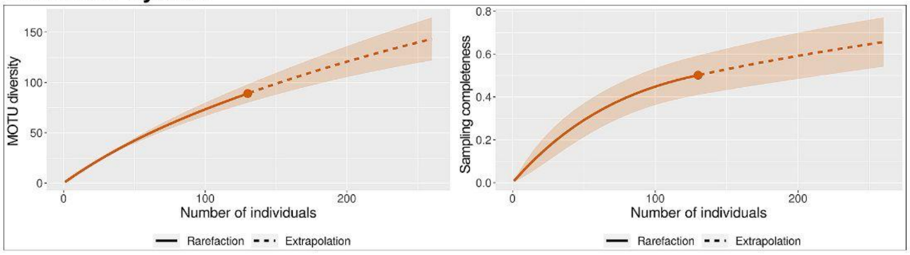


#### Sunda Shelf

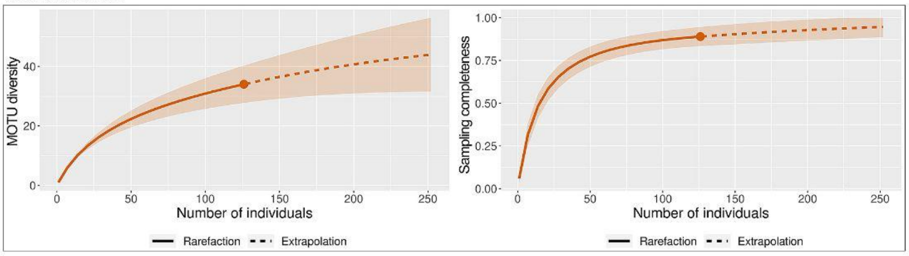


#### Tropical Southwestern Pacific

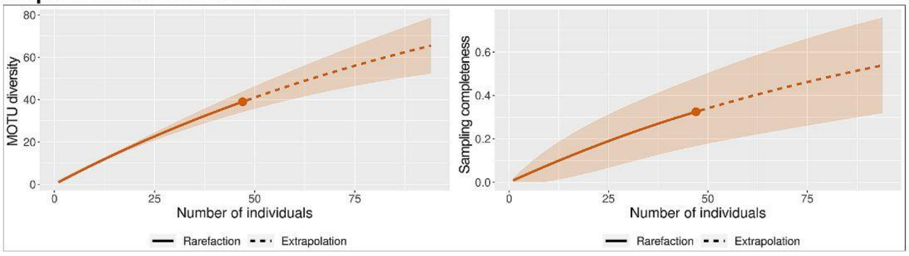


#### Western Indian Ocean

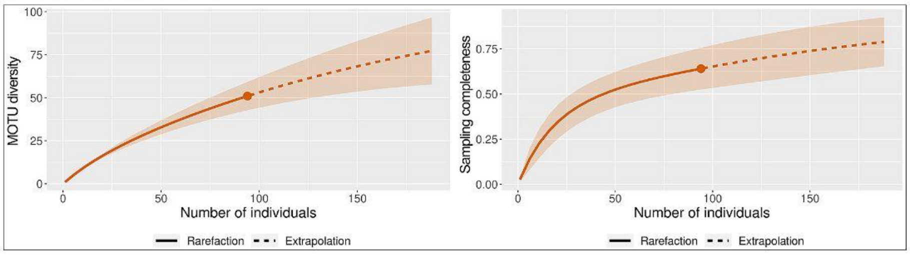
