## Appendix S3 for "Barcoding-inferred biodiversity of shallow-water Indo-Pacific demosponges"

**Figure S3.1:** 28S Jaccard dissimilarity for sponges of the studied marine provinces with singletons excluded. Cell colour also corresponds to the legend depicted in Figure 3, plus black in case of complete dissimilarity (100).

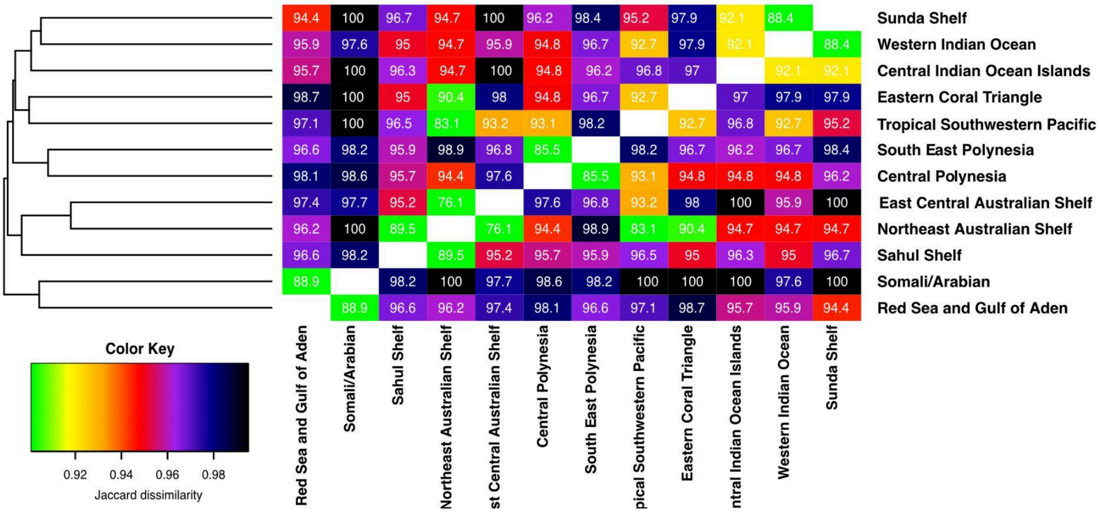

**Appendix S3 (continued)**

**Figure S3.2:** 28S Sørensen dissimilarity for sponges of the studied marine provinces including singletons. Cell colour also corresponds to the legend depicted in Figure 3, plus black in case of complete dissimilarity (100).

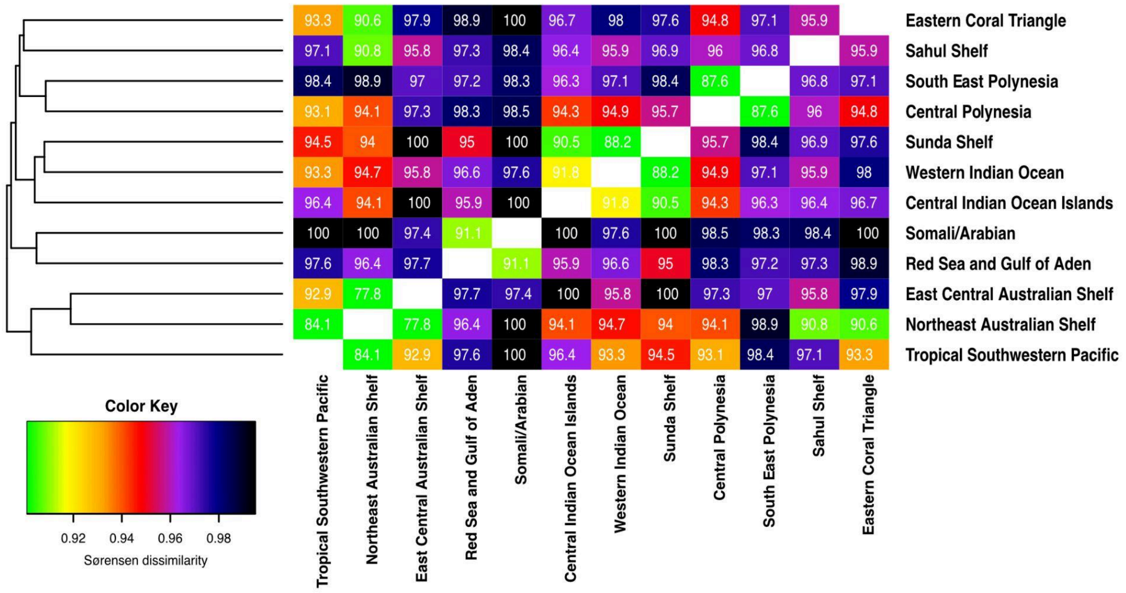

**Appendix S3 (continued)**

**Figure S3.3:** 28S Sørensen dissimilarity for sponges of the studied marine provinces with singletons excluded. Cell colour also corresponds to the legend depicted in Figure 3, plus black in case of complete dissimilarity (100).

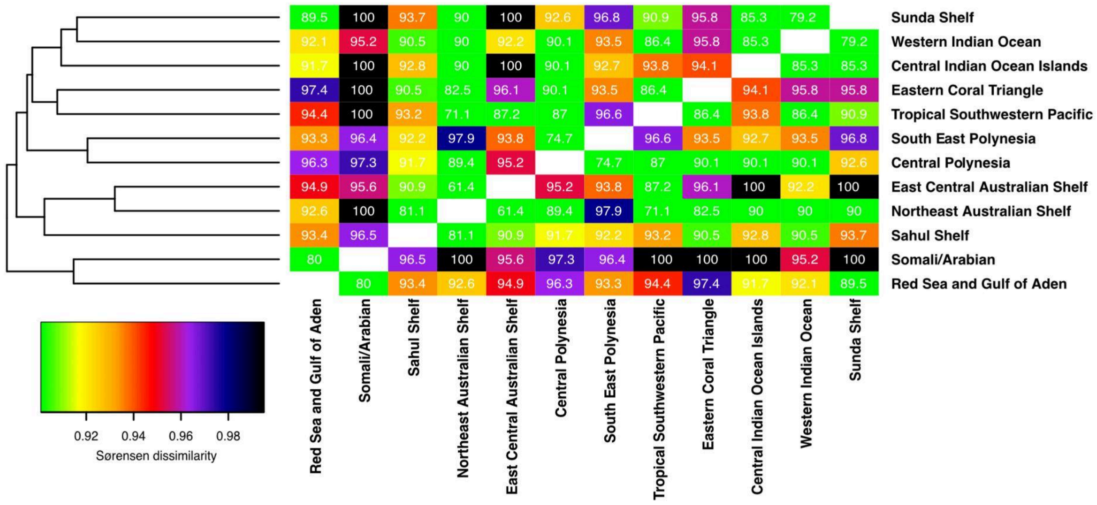
