## Appendix S4 for "Barcoding-inferred biodiversity of shallow-water Indo-Pacific demosponges"

**Figure S4.1:** Overview across marine province sequence yield and MOTU numbers in the analysis of faunistic boundaries for sponges from the *Central Indo Pacific* (yellow) and *Temperate Australasia* (blue) marine provinces around Australia.

| Realm (cf. Spalding et al 2007) | Marine Province | Seqs. total | Seqs. included | MOTUs | endemic MOTUs |
|---|---|---|---|---|---|
| ***Central Indo Pacific*** | ***Tropical Southwestern Pacific*** | 57 | 45 | 36 | 69.4% |
| | ***Northeast Australian Shelf*** | 207 | 181 | 96 | 68.8% |
| | ***Sahul Shelf*** | 166 | 151 | 96 | 89.6% |
| ***Temperate Australasia*** | ***East Central Australian Shelf*** | 79 | 70 | 44 | 61.4% |
| | ***Southeast Australian Shelf*** | 31 | 26 | 22 | 90.9% |

**Figure S4.2:** Jaccard dissimilarities between *Central Indo Pacific* (yellow) and *Temperate Australasia* (blue) marine provinces around Australia. Values shaded in green and red are discussed in the main text of the publication.

| Marine Province | ***Tropical Southwestern Pacific*** | ***Northeast Australian Shelf*** | ***East Central Australian Shelf*** | ***Southeast Australian Shelf*** |
|---|---|---|---|---|
| ***Sahul Shelf*** | **98.5** | **95,1** | **97.8** | **100** |
| ***Tropical Southwestern Pacific*** | | **91.8** | **96.1** | **100** |
| ***Northeast Australian Shelf*** | | | **88.0** | **98.3** |
| ***East Central Australian Shelf*** | | | | **98.4** |
